# Widespread and Abundant CRISPR-Cas Systems in the Deep Ocean

**DOI:** 10.1101/2025.05.26.656144

**Authors:** Pablo Sánchez, Marta Ferri-Peradalta, Ester María López-García, Raúl Ruiz, Belén Esquerrà Ruvira, Josep M. Gasol, Dolors Vaqué, Takashi Gojobori, Susana Agustí, Carlos M. Duarte, Francisco J. M. Mojica, Julián Cerón, Felipe H. Coutinho, Silvia G. Acinas

**Author notes:** These authors contributed equally (Pablo Sánchez, Marta Ferri-Peradalta).

## Abstract

**Background:** CRISPR-Cas systems have revolutionized modern biology. Most CRISPR-Cas systems used for biotechnological applications derive from cultivated bacteria, but evidence suggests that environmental microbiomes harbor an untapped diversity of these systems. Understanding of which environmental and biological factors drive the prevalence of CRISPR-Cas systems remains limited.

**Results:** A search for CRISPR-Cas systems was conducted among 176 globally-distributed marine metagenomes from the Malaspina expedition, including free-living and particle-attached microbiomes with emphasis on the deep ocean. CRISPR-Cas systems were more abundant in the deep ocean than in the surface and among free-living compared to particle-attached communities, reflecting the higher abundance of archaea and their viruses in these habitats. We identified 1,146 CRISPR-*cas* loci, some displaying unique architectures. A total of 48 Cas9 proteins were identified, many of which are potentially novel.

**Conclusions:** These discoveries expand the diversity of CRISPR-Cas and point at the deep-sea as a reservoir for future bioprospecting efforts of these resources.

## Background

Since their discovery, CRISPR-Cas (Clustered Regularly Interspaced Short Palindromic Repeats – CRISPR associated proteins) systems have grown to be a powerful and widespread biotechnological tool for genome editing, bringing forth enormous advances in the fields of molecular biology and genetics, with a growing potential for medical and pharmaceutical applications^1,2^. CRISPR-Cas are one of the many molecular defense systems that Bacteria and Archaea^3,4^ use to fend off invasive mobile genetic elements. CRISPR-Cas are unique among those in that they provide adaptive immunity, a mechanism which was previously believed to only be present in vertebrates. The system works by incorporating short fragments of invasive DNA as spacers between repeats within the CRISPR array, which then guides Cas proteins into cleaving invasive DNA or RNA matching such spacers. This allows the cell to remember, recognize, and clear future infections from mobile genetic elements that had previously been in contact with the cell. CRISPR were first observed fortuitously in *Escherichia coli*^5^ and more than a decade later, they were formally named and characterized in Archaea from hypersaline ecosystems^6,7^. In fact, higher prevalence of CRISPR-Cas systems have been observed among genomes of Archaea than Bacteria, as well as among extremophilic rather than among mesophilic prokaryotes^8^.

CRISPR loci are composed of several noncontiguous direct repeats separated by stretches of variable sequences, the spacers (often derived from viruses or plasmids) are commonly located next to a *cas* locus^9^. Those genes encode a large and heterogeneous repertoire of proteins, many of which interact with nucleic acids through domains that are homologues to those of nucleases, helicases, polymerases, and polynucleotide-binding proteins^10^. CRISPR-Cas systems can be classified according to the architecture of the CRISPR-Cas genomic loci, alongside the composition of homologous genes that make up the interference module. Currently, CRISPR-Cas systems are classified into two major classes, seven types, and a growing number of subtypes^11–16^. Class 2 systems are of particular interest since their effector complexes are composed of a single protein such as Cas9 or Cas12, which are the most commonly used for CRISPR-Cas genome editing applications, particularly Cas9^17^.

The ever-growing number CRISPR-Cas system discoveries in uncultured Bacteria and Archaea from environmental datasets^8,18–21^, and our new understanding of the extension of their genomic diversity^22–24^, suggest that environmental microbiomes are likely a major source of yet undiscovered CRISPR-Cas systems^8^. The oceans are the largest biome on Earth and encompass a large diversity of microorganisms resulting from over 3.5 billion years of evolution^25^. The abundances, taxonomic composition, and genetic repertoire of marine microbiomes shift in response to environmental gradients^3–25^, and also regarding their preferred habitats, e.g. free-living (FL), particle-attached (PA), or switching between both^29–32^. A recent study analyzing more than 40,000 bacterial and archaeal Metagenome Assembled Genomes (MAGs) from the Global Ocean Microbiome Gene Catalogue (GOMC) led to the identification of 5,127 diverse *cas* operons across 3,212 MAGs, amounting to a prevalence of about 15%^8^. The same study identified a novel CRISPR–Cas system harboring a Cas9 nuclease (Om1Cas9) with strong *in vitro* genome editing performance^8^.

Deep ocean microbiomes face specific constraints related to darkness, low temperature, and high pressure, which drive their community composition^33^, trophic strategies^27^, and virus-host interactions^34^. However, the factors influencing the prevalence and diversity of CRISPR-Cas in deep ocean ecosystems remains poorly characterized. Hence, we hypothesized that: i) the unique conditions of deep ocean ecosystems influence the prevalence of CRISPR-Cas systems, and the systems therein likely function at physiological conditions different from those of CRISPR-cas systems currently used for biotechnological applications, and ii) that poorly characterized taxa of deep ocean Bacteria and Archaea are untapped sources of novel CRISPR-Cas systems^8,35^. We test these hypotheses through the analysis of CRISPR-Cas elements among 176 globally-distributed marine microbial metagenomes retrieved by the Malaspina expedition^36^, which sampled both free-living and particle-attached microbiomes with emphasis on the deep ocean.

## Results

### Identification and classification of cas loci

We used 176 global ocean microbial metagenomes collected during the Malaspina Expedition^27,36,37^ (**Supplementary Table 1**) to characterize the abundance and diversity of CRISPR-Cas systems throughout broad environmental and habitat gradients of the pelagic global ocean. This dataset harbors a unique genetic repertoire not widely represented in other public microbiome gene catalogs and MAG databases^27^. These metagenomes include: i) a wide geographical distribution covering tropical and subtropical regions (**Figure 1A**), ii) depth-resolved vertical profiles spanning photic and aphotic zones with emphasis on bathypelagic habitats, which are underexplored ecosystems, iii) sampling of multiple plankton size fractions, iv) associated environmental metadata.

**Figure 1.**
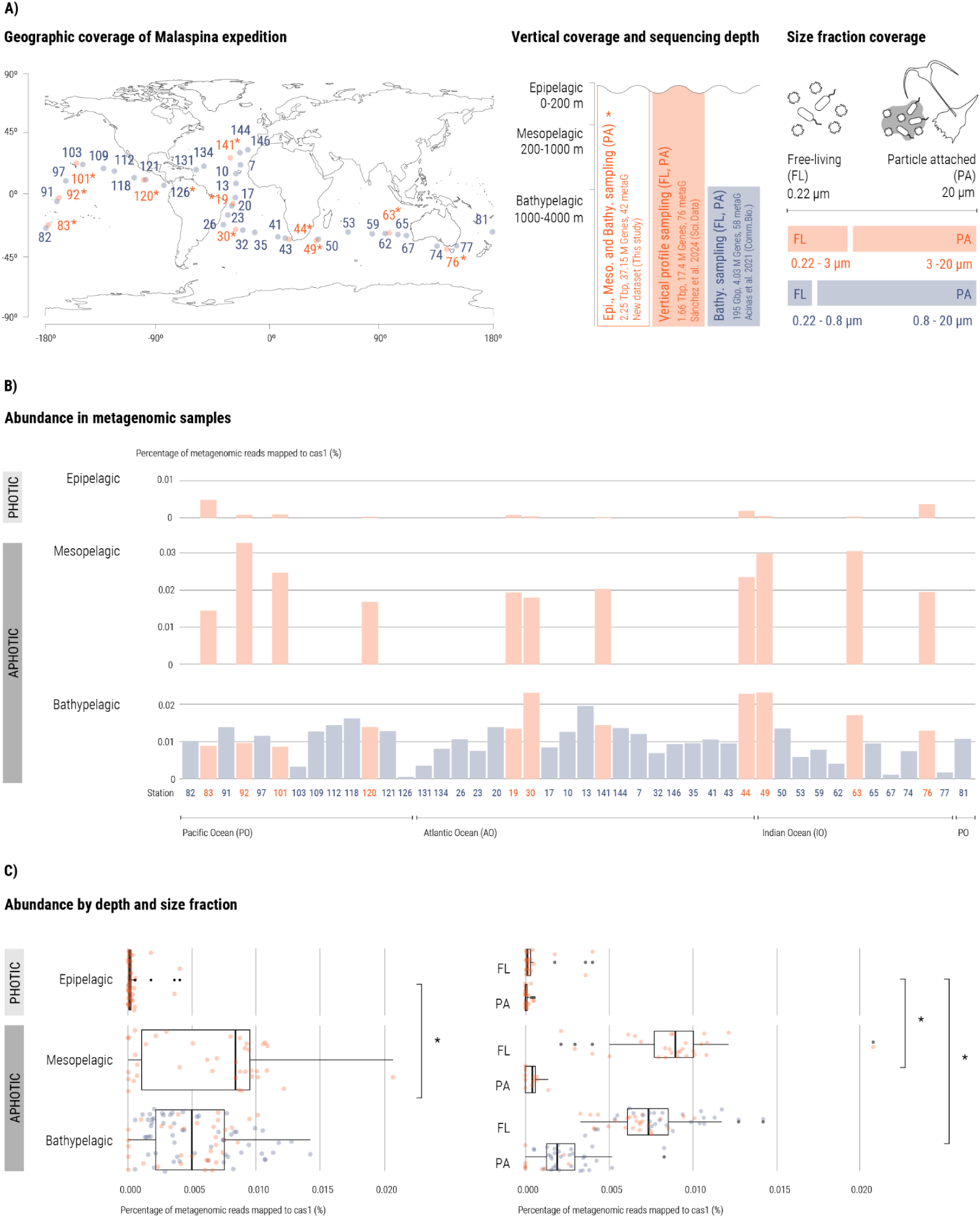
Malaspina expedition metagenomic sampling context and abundance of cas1 genes in metagenomic samples. (A) Left panel: geographic location of sampling sites. Sites depicted in orange are those where vertical profiles were obtained, while those depicted in blue are sites where only the bathypelagic was sampled. Middle panel: water column (vertical) and sequencing coverage of metagenomic samples. Right panel: size fraction representation of metagenomic samples used in this study. Metagenomic samples are classified into three batches: i) vertical profile metagenomes (Sánchez et al., 2024) cover epi-, meso- and bathypelagic layers, considering the free-living (FL) plankton size fraction between 0.22-3 μm and the particle attached (PA) size fraction between 3-20 μm; ii) bathypelagic metagenomes (Acinas et al., 2021) consider the FL plankton size fraction between 0.22-0.8 um and the PA size fraction between 0.8-20 μm. (B) Abundance of cas1 genes in metagenomic samples, represented as the number of reads corresponding to the cas1 gene relative to the total metagenomic reads in each sample. (C) Abundance of cas1 genes (as in panel B) categorized by water column layer (left panel) and size fraction (right panel). Asterisks indicate significant *p*-values (< 0.01).

We screened 98.2 million coding DNA sequences predicted among the assembled Malaspina metagenomes, leading to the identification of 528,589 coding DNA sequences that matched at least one of the 437 custom Hidden Markov Models (HMM) Cas protein profiles that made up our reference database. Out of these, only 4,707 proteins were finally classified as part of a CRISPR-Cas locus (**Supplementary Table 2**). In total, we detected 1,146 CRISPR-Cas loci, 70% of which were classified unambiguously as defined by the majority rule approach (see methods), and 43% contained at least one effector protein and therefore were considered complete loci (**Supplementary Table 3**). Aiming to reduce the possibility of obtaining false-positives, further analyses were restricted to those CRISPR-Cas loci containing at least three genes encoding Cas proteins. However, this is also likely to lead to overlooking some true-positives, especially some class 2 systems, as well as some variants of types V and VI systems^38,39^. The majority of putative CRISPR-Cas loci were assigned to type I systems (**Supplementary Figure 1**). The higher prevalence of class 1 systems was also observed in other environmental microbiome studies, including extreme environments such as hot spring microbial communities^8,19,40^.

### Drivers of abundance and prevalence of CRISPR-cas systems across the global ocean

We used the relative abundances of Cas1 within metagenomes as a proxy for the relative abundance of CRISPR-Cas systems in the samples, as Cas1 is ubiquitous among CRISPR-Cas systems capable of autonomous spacer acquisition. The microbial metagenomes from Malaspina Expedition aimed at covering the range of cell-sizes and lifestyles (i.e. free-living or particle-attached) among planktonic communities of bacteria, archaea, and microeukaryotes. Thus, the dataset included metagenomes from different plankton size-fractions, i.e. 0.22-0.8 μm, 0.22-3 μm, 0.8-20 μm, and 3-20 μm (**Figure 1A**). The relative abundance of CRISPR-Cas systems was significantly higher in the aphotic ocean (i.e. the mesopelagic and bathypelagic) than in the photic zone (epipelagic) (**Figures 1B and 1C)**. Among the aphotic zone samples CRISPR-Cas relative abundances were significantly higher among free-living (FL) than particle-attached (PA) samples, while no significant differences between these metagenomes representing these lifestyles were observed among photic zone samples (**Figure 1C**). To further validate these findings we performed a complementary analysis in which, instead of using Cas1 protein from the Malaspina dataset, we quantified all the Cas proteins identified by CRISPRCasTyper^41^ in MAGs from the OceanDNA^24^ database (**Supplementary Figure 3**). We observed consistent enrichment patterns when comparing aphotic and aphotic free-living fraction samples from the vertical profiles dataset *(p*-value = 0.013), and when comparing free-living and particle-attached samples from the size-fractioned bathypelagic dataset (*p*-value = 0.28). The fact these patterns were recovered using a different tool for identifying CRISPR-cas systems, reference genomes, and database for quantifying these systems, provides evidence that these findings did not emerge from biases associated with our methodology for identifying and quantifying Cas proteins in the metagenomes in the first place.

Higher temperatures have been observed to be associated with an increase of CRISPR-Cas, driven by the stark differences in the prevalence of CRISPR-Cas systems among genomes of psychrotrophs (1%), mesophiles (11%), and thermophiles (40%)^8^. Our findings suggest that the cold temperatures of the deep ocean, namely 2.0 ± 0.5 °C in the Malaspina bathypelagic samples (3,731 ± 495 m deep)^27^, and 2.6 ± 1.3 °C in the Malaspina vertical profiles samples from 1,000 to 4,000 m deep^36^, represents an extreme biome with higher abundances of CRISPR-Cas systems.

We hypothesized that the observed patterns of CRISPR-Cas abundances derive from the marked differences on the taxonomic composition of microbial communities observed between photic and aphotic layers^28,42^, as well as those between free-living and particle-attached habitats^27,32,34,43,44^. Namely, free-living bathypelagic communities are characterized by oligotrophic, slow-growing generalists living in low cell density environments. Meanwhile, their particle-attached counterparts are characterized by copiotrophic, fast-growing specialists which grow quickly while nutrient availability is high^27,32^. We tested this hypothesis by comparing the relative abundances of OceanDNA MAGs between photic and aphotic samples, and between free-living and particle-attached samples. A total of 2,888 genomes were found to be significantly more abundant among photic samples, while 3,332 were found to be significantly more abundant among aphotic samples, with marked differences in the taxonomic affiliations of each group (**Supplementary Figure 4A**). Next we compared the number of identified cas operons per genome between the genomes of each group, which was significantly more abundant among those enriched in the aphotic zone (*p*-value = 2e-13). This corroborated the patterns observed when quantifying the relative abundances of Cas proteins (**Figure 1** and **Supplementary Figure 3**). Meanwhile, a total of 3,327 genomes were found to be significantly more abundant among bathypelagic free-living samples, while 824 were found to be significantly more abundant among bathypelagic particle-attached samples (**Supplementary Figure 4B**). In this case, the observed pattern contradicted the results observed when quantifying the relative abundances of Cas proteins (**Figure 1** and **Supplementary Figure 3**), as the number of cas operons detected among among genomes enriched in particle-attached samples was significantly higher (*p*-value < 2e-16). This discrepancy could be attributed to an underrepresentation of particle-attached genomes in the OceanDNA databases as these organisms tend to be less abundant than their FL counterparts, and few studies enrich this particular size-fraction when preparing metagenomes.

We next sought to determine if the higher CRISPR-Cas abundances in the aphotic zone, and in the FL samples from this layer were associated with the abundances of Archaea, as archaeal genomes display higher prevalence of CRISPR-Cas systems than bacterial genomes^8,35^. Archaea can amount to as much as 20 % of all prokaryotic cells in the deep ocean^45–47^. Among the metagenomes from the Malaspina dataset retrieved across depth and geographical gradients (0.22-3 μm size fraction), Archaea belonging to phylum Thermoproteota were the second most abundant taxa among mesopelagic samples^37^. In the 58 size-fractioned bathypelagic metagenomes, Archaea accounted for 21% of the reads and their relative abundances were higher in the FL fraction samples than in their PA counterparts^4^. Thus, we analyzed the relative abundances of Archaea and Cas 1 proteins in the 76 metagenomes of the FL size fraction from the Malaspina dataset covering depth gradients. This revealed a strong positive association between these two variables (Spearman’s rank correlation coefficient, ⍴ = 0.89), driven by their high abundances in the deep ocean, especially in the mesopelagic zone (**Figure 2A**).

**Figure 2:**
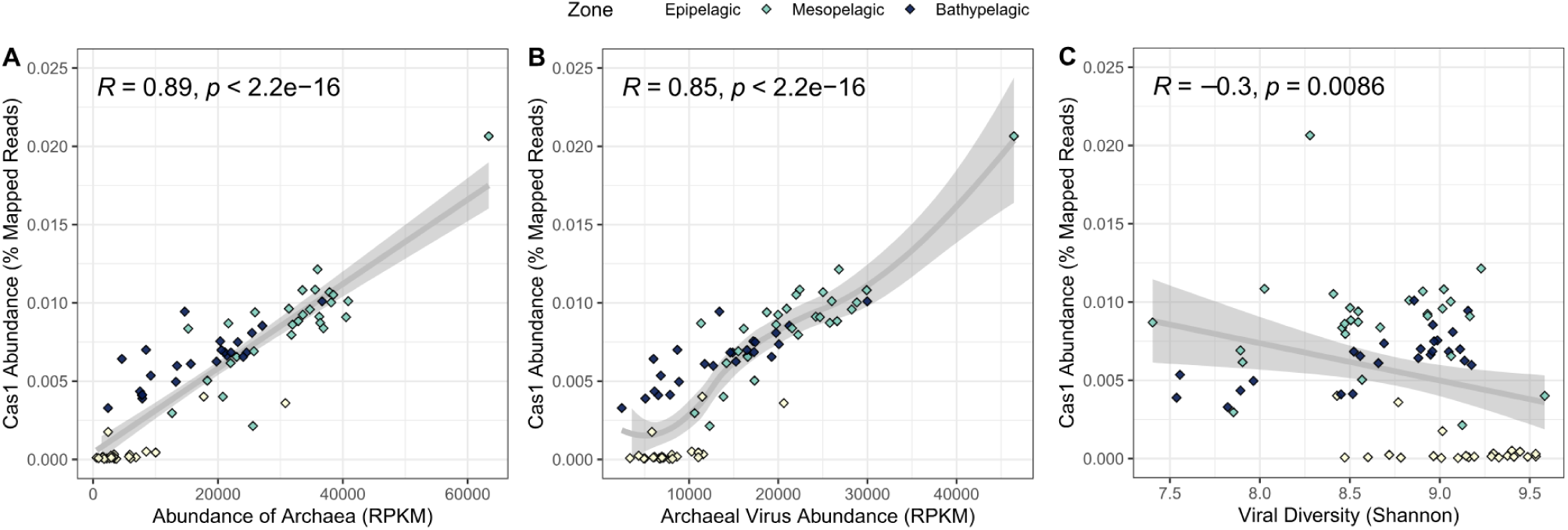
Association between Cas1 abundances and metrics of community composition in 76 Malaspina depth profiles metagenomes from the 0.22-3 μm size-fraction. In all scatterplots the y-axis depicts the relative abundances of CRISPR-Cas in the metagenomes. Samples are coloured by depth zone. Gray lines depict regression model fits between variables with 0.95 confidence intervals depicted by shaded areas. Spearman correlation coefficients and associated *p*-values are displayed in the upper left corners. A) Association with the relative abundance of Archaea. B) Association with the relative abundance of Archeal viruses. C) Association with the Shannon diversity index of the viral communities.

High viral abundance and low diversity were associated with increased CRISPR-Cas prevalence across diverse ecosystems^35^. In our study, the association between viral community diversity and CRISPR-Cas abundances yielded a weaker, yet significant negative association (⍴ = −0.3, **Figure 2B**). Analyzing the association with the relative abundances of viruses predicted to infect Archaea led to a similar positive correlation (⍴ = 0.85) to that observed for their hosts (**Figure 2C**). Interestingly, these correlations were much weaker when considering viruses of Bacteria, and not significant when all viral sequences were considered together (**Supplementary Figure 2**). Notably, when these correlations were calculated separately for each depth zone, the positive associations between relative abundances of CRISPR-Cas and viruses of archaea were also identified, although they were weaker among epipelagic subset (⍴ = 0.47; *p*-value = 0.023) than in mesopelagic (⍴ = 0.79; *p*-value = 2.2e−06) and bathypelagic (⍴ = 0.8; *p*-value = 4.8e−06) subsets. These results align with previous findings that linked CRISPR-Cas abundance with reduced prokaryotic cell abundances^48^ and lower viral diversity^35^. Our data expands these findings by demonstrating how CRISPR-Cas abundance shifts across environmental gradients among different ocean layers and among free-living and particle-attached microbial niches in the deep ocean, and how it is related to the abundance of archaea and the abundance and diversity of the viruses that infect them.

### CRISPR Cas9 loci architectures

We focused our interest in CRISPR-Cas9 loci due to their relevant use in genome editing as single effectors of class 2 systems^17^. We scanned 1,146 CRISPR-Cas loci, and obtained a collection of 48 Cas9 proteins (**Supplementary File 1** and **Supplementary Table 4**). Interestingly, we found that 62% of Cas9 encoded in our dataset (30 out of 48) were on average 194 amino acids shorter than SpCas9, *Streptococcus pyogenes*’ Cas9, the most widely used protein for genome editing (1,174 ± 377 amino acids vs. 1,368 amino acids respectively), suggesting that this smaller size could facilitate the use of these proteins in delivery systems.

Multiple *cas* loci architectures were identified (**Figure 3**). The most frequently observed loci architecture was *cas1*, *cas2* and *cas9,* which accounted for 25 of the 48 total loci. These CRISPR-Cas9 systems were distributed among six different orders of bacteria, such as Gammaproteobacteria, Alphaproteobacteria, or Flavobacteria. This CRISPR locus composition was observed across all plankton size fractions (FL and PA), depths, and ocean regions. Among the other loci architecture identified, we found those in which genes encoding for ancillary proteins (CasR) were present, which were always associated with genomes of the order Flavobacteriales, and loci in which the Cas2 was replaced by a DEDDh exonuclease, as well as unusually short Cas9 proteins (one of them only 378 amino acids long), found in association with Cas4 and DEDDh encoding genes (**Figure 3**). We found another putative novel locus formed by the type-II miniature effector protein Cas14 with Cas9, Cas1, and Cas2, which was only described previously in a locus with Cas1, Cas2, and Cas4^49^. This locus was found in association with four bacterial orders (Cytophagales, Flavobacteriales, Balneolales and Sphingomonadales), and it was found only in the FL fraction of the aphotic layers (mesopelagic and bathypelagic) from the Atlantic Ocean. The finding of such loci with a small effector protein next to a full-sized Cas9 raises the hypothesis of whether these shorter effector versions are functional, and if they could play a complementary role.

**Figure 3.**
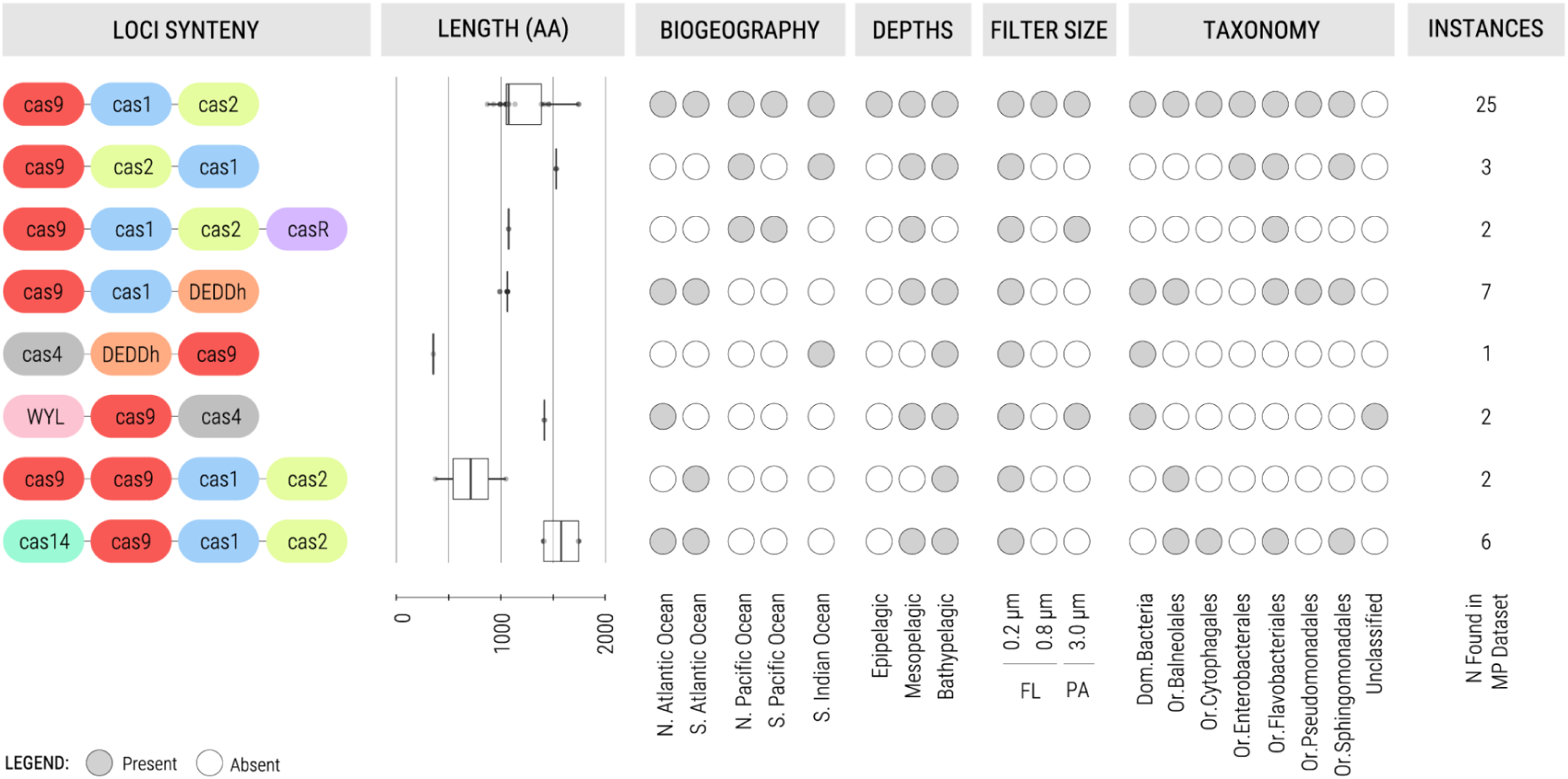
Characterization of CRISPR-Cas loci in the Malaspina expedition metagenomes. Loci synteny of cas systems in the Malaspina expedition is followed by the characterization of their Cas9 protein length (in boxplot format), biogeographic distribution, vertical profiling (depths), filter size (FL: free-living and PA: particle-attached fractions), taxonomic classification and the number of observations (instances) found in the dataset. Filled circles represent presence, empty circles represent absence.

In terms of geographical patterns of diversity, station 44 in the South Atlantic Ocean contained the most diverse range of loci architectures (**Supplementary Figure 1**), particularly concentrated in the bathypelagic layer. This location corresponds to the Agulhas Current, one of the major oceanic currents, that connects the Indian, Atlantic and Southern Oceans^50^. This elevated diversity of loci architectures observed here may be due to this unique feature, as currents could drive microorganisms harboring different types of CRISPR-Cas systems initially located far from each other across different oceans to this specific area.

None of the CRISPR-*cas9* loci were assigned to archaea, as these organisms seldom harbor such systems in their genomes^51^. Also, the CRISPR-Cas 9 type II-A (to which the canonical SpCas9 belongs to), was absent in our study as it is usually rare among environmental microbiome ^8,40^. Although we overlooked the identification of potential CRISPR-Cas9 in mobile genetic elements such as plasmids^52^ in our dataset, it seems that the II-A subtype is overrepresented in reference databases due to its high prevalence among human-associated pathogenic bacteria, but those are not representative of those found among the ocean microbiome.

### Taxonomic annotation and phylogenetic reconstruction of Cas9 proteins

We determined the likely organisms from which the identified Cas9 proteins were derived. To that end, we queried the 48 Cas9 proteins against a database of proteins derived from MAGs from the Malaspina catalog^27,37^, and from the OceanDNA database^24^ through best-hit classification. All 48 Cas9 proteins matched homologues derived from MAGs, out of which 36 yielded matches with 100% identity (**Supplementary Table 4**). The identity levels from the 12 remaining proteins ranged from 99.9% to 25.9% (mean = 57%). A total of six Cas9 sequences displayed less than 50% amino acid identity. In terms of taxon prevalence, Cas9 proteins were most often linked to MAGs classified as Bacteroidota (23), Gammaproteobacteria (11), and Alphaproteobacteria (8). Notably, one Cas9 was linked to the phylum Marinisomatota, which was not previously reported as a potential source of CRISPR-Cas systems. In addition, many of the Cas9 proteins were linked to genera that are typically abundant in the bathypelagic, such as Erythrobacter_A, *Zunongwangia*, or *Leeuwenhoekiella*^53^, pointing to these taxa as potential underexplored reservoirs for yet undiscovered CRISPR-Cas systems.

A phylogenetic reconstruction revealed that most of Malaspina Cas9 proteins clustered with those from the subtype II-C systems in CasPEDIA database^13^ (**Figure 4**), specifically within clades VI, VIII, IX, and X^14^. Only two Cas9 clustered together with those from type II-B (clade I) systems. As mentioned above, we did not identify any II-A system, which may reflect an association of such subtype to pathogens or habitats and taxa not abundant in marine microbiomes. Interestingly, not all Cas9 proteins derived from loci with the same architecture were grouped in the same phylogenetic cluster or assigned to the same taxa (**Figures 3 and Figure 4**), further highlighting the diversity of these systems within our dataset. Notably, many proteins described in the present work formed unique clades within the tree, often with long branch lengths, corroborating our hypothesis that the deep ocean is an overlooked reservoir, rich in novel Cas9 proteins with unexplored biotechnological potential.

**Figure 4.**
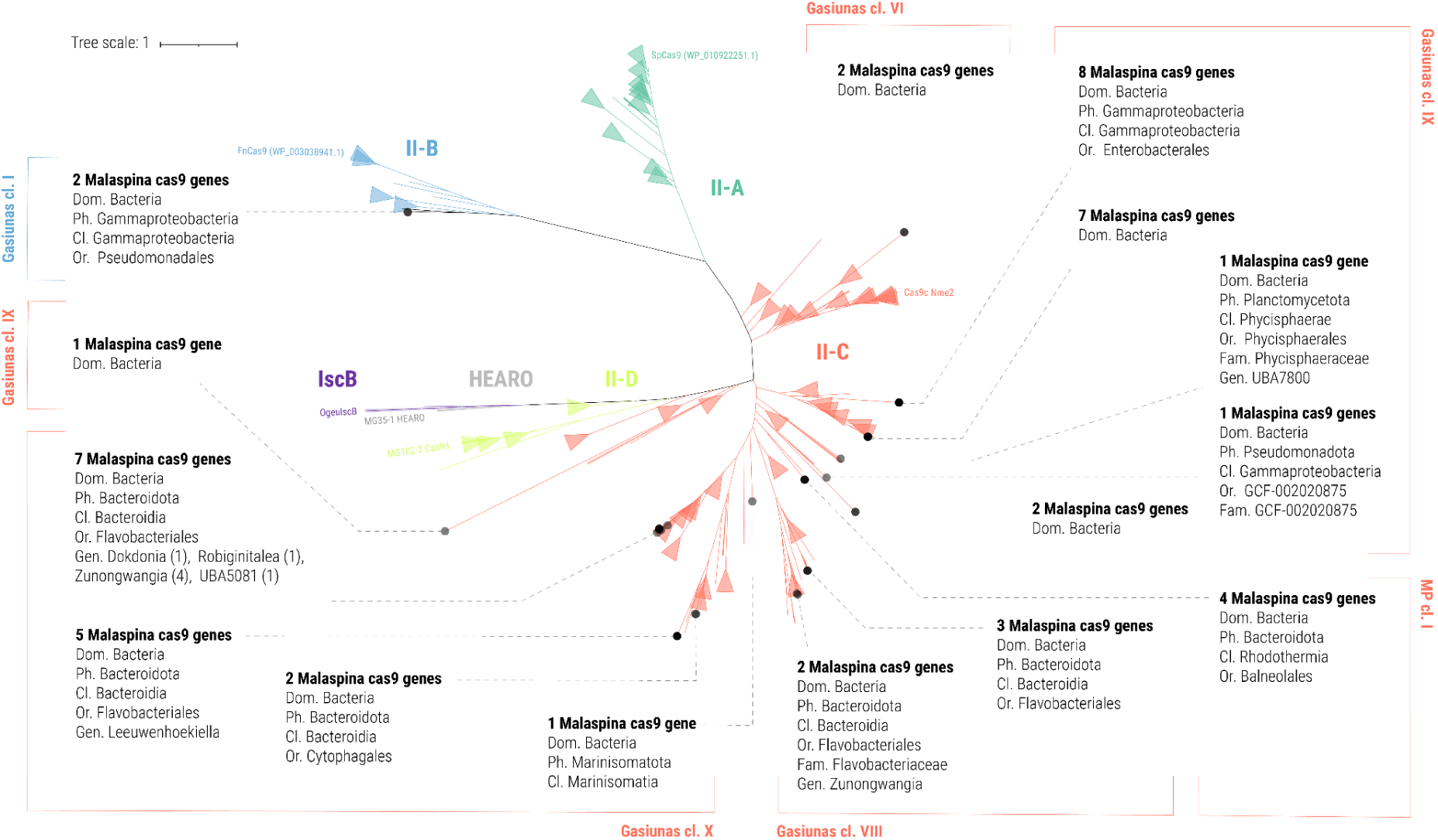
Phylogenetic tree of 48 Cas9 proteins found in the Malaspina expedition metagenomes. A phylogenetic tree was reconstructed by combining the 48 Cas9 proteins from the Malaspina dataset and those described in the CasPEDIA database to determine the phylogenetic placement of these Cas9 in light of previously characterized proteins. Tree branches are coloured based on the Makarova classification. Malaspina Cas9 proteins (black dots) are grouped according to their phylogenetic placement and taxonomic annotation and they are associated to a Gasiunas class in the outer brackets annotation. The taxonomic annotation of Malaspina Cas9 proteins was determined by best hit classification after querying them against a database of proteins from metagenome assembled genomes according to their degree of identity, i.e. Malaspina Cas9 proteins with best hits with identity ≥ 70% were assigned the same taxonomic annotation of the corresponding MAG to the deepest level, those with identity between 50% and 70% were assigned taxonomic annotation at the level of order, while those with identity below 50% were assigned taxonomic annotation at the level of domain. The cluster of 4 Balneolales Cas9 proteins (Makarova II-C) that could not be assigned to any known Gasiunas class is classified into new MP class I. Taxonomic annotation is detailed until the last known level.

### Conclusions

Altogether, these findings corroborate our hypothesis that deep ocean ecosystems are hot-spots of CRISPR-Cas system abundance and diversity. Furthermore, we provide evidence that this trend is linked to the abundance of Archaea and their associated viruses, that characterize the free-living microbial communities of the deep ocean, especially in the mesopelagic zone. Moreover, we demonstrate that poorly characterized marine bacterial taxa encode CRISPR-Cas systems in their genomes with atypical loci architecture harboring Cas9 proteins that belong to previously uncharacterized evolutionary lineages. These findings provide strong motivation to guide future bioprospecting efforts targeting marine ecosystems, with a particular attention to the deep sea, complemented with further experimental validation and characterization of the biochemical properties of these proteins. CRISPR-Cas9 systems retrieved from the deep sea will likely display optimal activities at specific environmental conditions, which would expand the use of CRISPR-Cas in biotechnology applications. This will provide further value to the applicability of deep sea Cas nucleases across different CRISPR-based applications.

## Methods

### Dataset description and analysis

A total of 176 metagenomes from the Malaspina Expedition 2010^36^ were analyzed derived from 43 stations, including 11 vertical profiles with 7 depths from surface to 4000 meters deep each with a total of 76 metagenomes from the FL size fraction (0.22-3.0 µm)^37^, 58 metagenomes from 32 stations characteristic from deep ocean water masses from the FL and PA size fractions (0.22-0.8 µm and 0.8-3.0 µm respectively)^27^ and 43 new nanoplankton metagenomes (3-20 µm) from 14 stations that cover 6 vertical profiles sampled at 3-7 depths, from the surface to 4,000 meters deep (or to the sea floor in shallower waters), plus 12 additional metagenomes. With these datasets we recovered a wide representation of the diversity of both free-living and particle-attached microbes of the global ocean, with emphasis on the deep ocean, totaling 4.12 Tbp of data (2.27 Tbp coming from the new metagenomes released here; 52.9 Gbp ± 16.0, mean ± standard deviation per sample, **Supplementary Table 5**). Metagenomic analysis was carried out as follows: sequencing reads were assembled with MegaHit v1.1.3 ^54^ with options “--presets meta-large --min-contig-len 500”. Genes were predicted from assembled contigs using Prodigal within Prokka v1.14.6 ^55^. Next, all genes were clustered at 95% of nucleotide sequence identity with CD-HIT-est v4.6.1 ^56^ and options “-c 0.95 -G 0 -aS 0.9 -g 1 -r 1 -d 0 -s 0.8” to obtain a non-redundant gene catalog of 51.5 million non-redundant genes, from which novel *cas* genes were mined.

### Detection and classification of Cas loci

A custom database was created by collecting Cas genes available from different studies with representation of all currently described types^18,19,38,39,49,57^. First, protein sequences were aligned with MAFFT v7.402^58^with default options. Hidden Markov Model (HMMs) profiles were built from the alignments using HMMER package v3.2 ^59^. Subsequently, Malaspina protein sequences were queried against the HMM profiles with hmmscan (e-value < 0.001). Finally, all the matches were parsed into GFF files and analyzed through a custom python v2.7.15 script designed to detect and classify Cas loci, using the approach that follows: a CRISPR locus was defined as a cluster of at least three consecutive Cas coding DNA sequences identified in the same contig, allowing for up to four non-*cas* genes in between them. Such loci were further classified into different CRISPR-Cas system (sub)types through a combination of two approaches. First, loci were assigned a system type classification following a majority rule when at least two thirds of the genes in the locus matched the HMM profiles on the individual proteins that make up a given system type. The second approach relied on the detection of maker Cas proteins that are specific from a particular (sub)type ^57^. The locus classification was considered unambiguous if the result of the majority rule was conclusive. Completeness of the CRISPR-Cas system was evaluated by the presence of effector proteins in the locus. We tested the performance of the detection and classification of Cas loci algorithm using a mock GFF file based on previously published data from classified loci ^57^. Further details on the algorithm and its validation can be found at the Gitlab repository of the project provided in the code availability section.

### Relative abundances and geographical distribution of CRISPR-Cas systems

To assess the presence and distribution of CRISPR-Cas systems across the global ocean, Cas1 proteins were used as a proxy to estimate relative abundances of CRISPR-Cas systems in Malaspina metagenomes^3^. We built a database containing the Cas1 proteins from the reference set used to build the HMM profiles, and those predicted in the Malaspina samples and we queried against the Illumina R1 reads from the metagenomes using MMSeqs2 (Development version, commit 15c7762) using parameters “-s 1.0 -e 0.00001 --min-aln-len 20 --min-seq-id 0.3 --max-seqs 5”. Relative Cas1 abundances in metagenomes were estimated as the sum of all unique reads mapping to Cas1 proteins divided by the total number of reads in the metagenome. For samples represented by multiple metagenomes sequenced in multiple batches, Cas1 abundances were defined as the median abundance across replicates. In addition, CRISPRCasTyper^41^ was used to identify CRISPR-Cas operon among over 8,000 species representative metagenome assembled genomes from the OceanDNA database^24^. All identified Cas proteins were pooled together for relative abundance quantification among Malaspina metagenomes as described above for the Cas1 proteins.

The relative abundances of bacterial, archaea, and viruses among metagenomes from the Malaspina vertical profiles were calculated as follows. Genomic sequences of bacteria, archaea, and viruses were obtained from a previously characterized dataset of metagenome assembled genomes and viral genomic sequences derived from Malaspina samples^37^. The taxonomic affiliation of the MAGs was assigned using GTDB-tk^60^ v1.7 Putative hosts of viral sequences were determined using PHIST^61^ v1.2.1. A reference database was built containing sequences from 1,228 de-replicated medium and high quality MAGs. A second reference database was built containing viral sequences. Next, post-QC reads from the metagenomes were queried against the reference databases using Bowtie2^62^ v2.3.4.1 in sensitive local mode. Output SAM files were converted to BAM and sorted using Samtools^63^ v1.15. Sequence relative abundances were calculated as Reads Per Kilobase per Million total sequences (RPKM). The abundances of archaea, bacteria, and viruses were calculated as the sum of RPKM values grouped according to taxonomic classification (for the MAGs) or predicted host taxon (for the viruses). The same strategy was used to quantify the relative abundances of the OceanDNA MAGs in the Malaspina metagenomes.

### Taxonomic annotation and phylogenetic reconstruction of Cas9 proteins

The 48 Cas9 proteins from the type II systems were subjected to best-hit classification by querying them through BLASTP (Minimum bitscore = 50 and maximum E-value = 0.001) against a dataset of proteins derived from marine MAGs from the OceanDNA database and Malaspina datasets^24,27,37^ which had previously been assigned taxonomic classification using GTDB-tk^60^. Taxonomic annotation of Malaspina Cas9 proteins was assigned according to their degree of identity, i.e. Malaspina Cas9 proteins with best hits with identity ≥ 70% were assigned the same taxonomic annotation of the corresponding MAG to the deepest level, those with identity between 50% and 70% were assigned taxonomic annotation at the level of order, while those with identity below 50% were assigned taxonomic annotation at the level of domain. For phylogenetic reconstruction, the Cas9 sequences obtained from the Malaspina dataset were combined with 830 Cas9 sequences from the CasPEDIA database^13^. Sequences were aligned with Muscle v5.1^64^ using the “-super5” parameter. Phylogenetic reconstruction was performed through IQ-TREE v2.0.6^65^ using the “-MFP” option, which identified VT+F+R10 as the best substitution model.

## Declarations

### Ethics approval and consent to participate

Not applicable

### Consent for publication

Not applicable

### Funding

The Malaspina 2010 Expedition was funded by the Spanish Ministry of Economy and Competitiveness (MINECO) through the Consolider-Ingenio program (ref. CSD2008-00077) to Carlos M. Duarte. Furthermore, this work has been supported by the grant Malaspinomics CTM2011-15461-E. Additional funding was provided by projects of the Plan Nacional I + D + I 2017 (CTM2017-87736-R), Deep Cas (PDC2021-120968-I00) to S.G.A from the Spanish Ministry of Economy and Competitiveness and PID2023-146919NB-C21 funded by MCIN/AEI/10.13039/501100011033/FEDER, UE to S.G.A and P.S from the Spanish Ministry of Science, Innovation and Universities. Members of the ICM had the institutional support of the “Severo Ochoa Centre of Excellence’’ accreditation (CEX2024-001494-S) funded by AEI10.13039/501100011033. F.J.M.M. is supported by grants PID2023-150750NB-I00 (funded by MICIU/AEI/ 10.13039/501100011033 and by ERDF/EU), PROMETEU/2021/057 (funded by Conselleria d’Educació, Cultura, Universitats i Ocupació, Generalitat Valenciana, Spain), and INNEST/2024/427 (funded by Agencia Valenciana de Innovación - IVACE+I Innovación - and by the European Union through the ERDF Program of the Valencian Community 2021-2027). The activity of J.C in this project is supported by the grant PID2023-146930NB-I00 from the Spanish Ministry of Science, Innovation and Universities. F.H.C was supported by a Ramón y Cajal fellowship (RYC2022-037094-I) from the Spanish Ministry of Science and Innovation.

### Availability of data and materials

The sequences, MSAs and HMMs profiles of the novel putative families, and the Cas9 sequences used to build the tree can be downloaded from the project website after publication. Malaspina metagenomes have been deposited in the European Nucleotide Archive under project PRJEB52452. All the code and other data used in the project can be obtained from the project Gitlab repository (https://gitlab.com/martaferri/malaspina.crispr-cas).

## Acknowledgements

We thank the R/V Hesperides crew, the chief scientists in Malaspina legs, and all project participants for their help in making this project possible. We thank the maintainers of the Marine Bioinformatics Core (MARBITS) of the Institut de Ciències del Mar (ICM-CSIC) where the High-Performance computing data analysis was performed. F.H.C. was granted access to the Galician Supercomputing Center (CESGA) infrastructure where some of the data analysis was performed. The supercomputer FinisTerrae III and its permanent data storage system have been funded by the Spanish Ministry of Science and Innovation, the Galician Government and the European Regional Development Fund (ERDF).

## Competing interests

The authors declare that two *cas9* patents derived from this dataset have been filed under numbers EP24383088 and EP24383089.

## Authors’ contributions

CMD,SGA, JMG, and DV designed and carried out the sampling and sample processing strategy. SGA,FHC, PS, FJMM, and JC designed data analysis pipelines. PS, MFP, EMLG, and FHC analysed the data. All authors contributed to data interpretation and to writing of the manuscript.

## Supplementary material

**Supplementary Figure 1:**
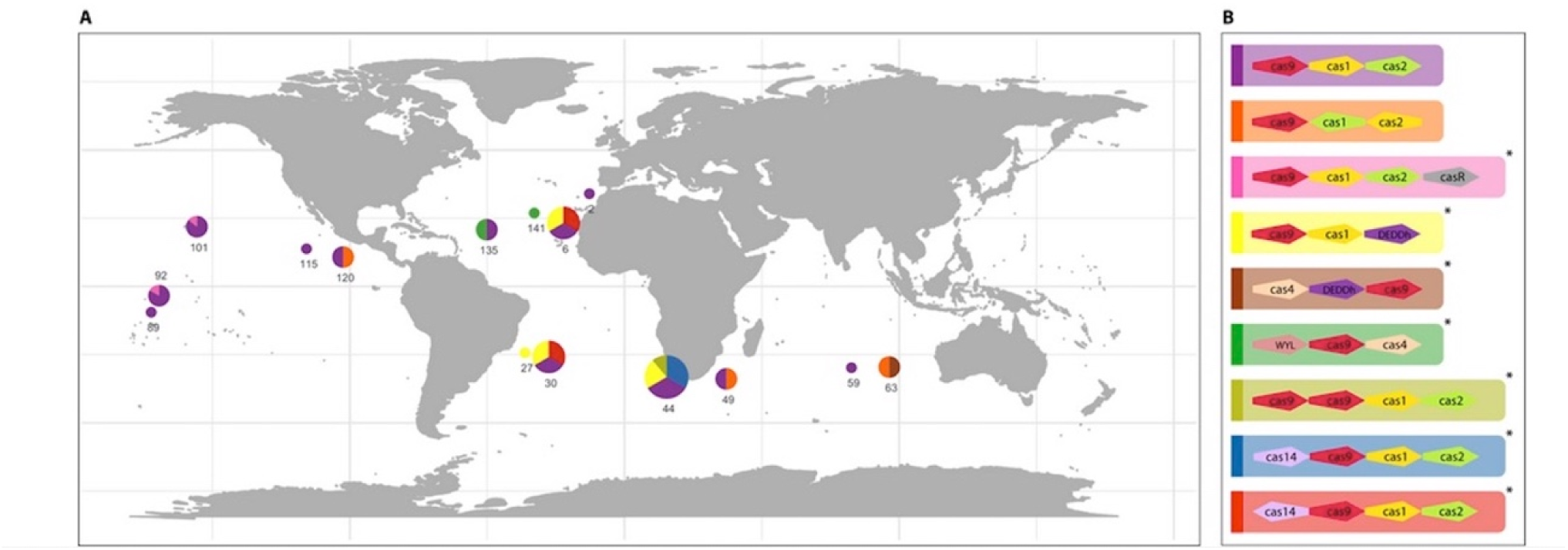
Diversity and prevalence of Cas9 containing loci architecture across sampling sites. Different colors represent different CRISPR-Cas9 loci Architectures from Figure 3 (*represent potential new loci).

**Supplementary Figure 2:**
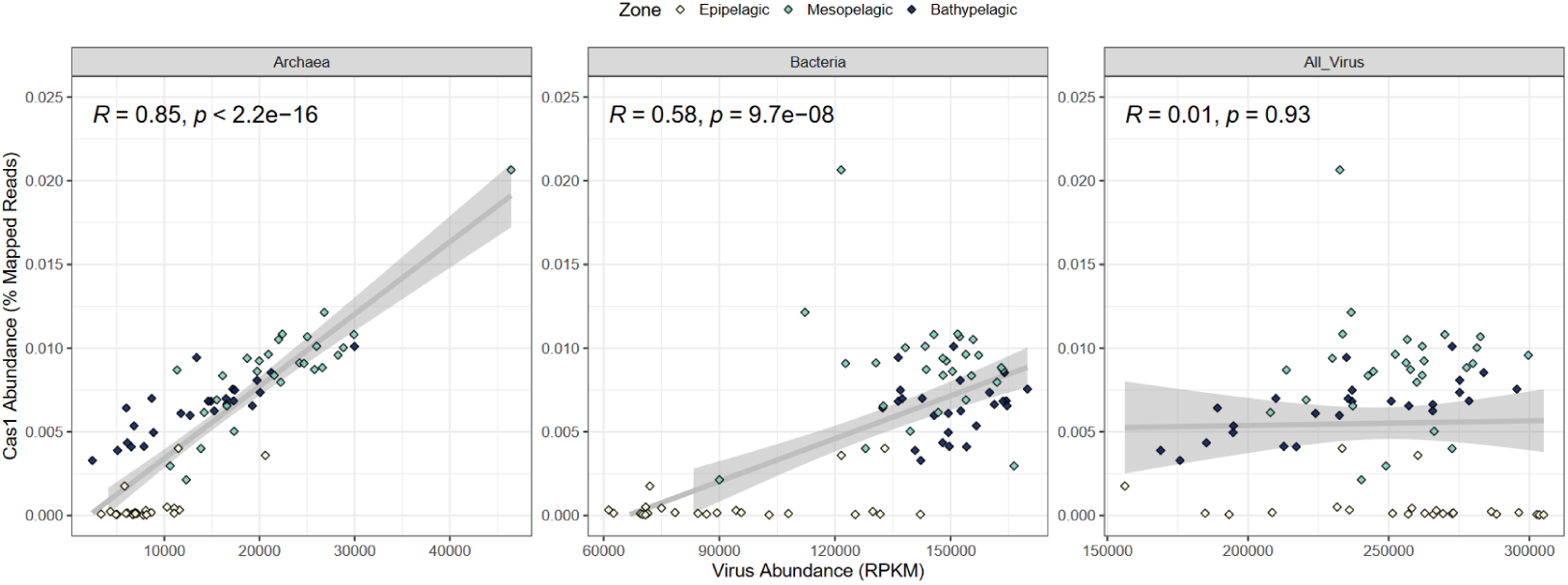
Association between relative abundances Cas1 and viruses in 76 Malaspina depth profiles metagenomes from the 0.22-3 μm size-fraction. The left and middle panels display the sums of relative abundances of viruses according to the predicted host domain. The right panel depicts the sum of relative abundances of all viral sequences, including those that could not be assigned a putative host. In all scatterplots the y-axis depicts the relative abundances of CRISPR-cas in the metagenomes. Samples are coloured by depth zone. Gray lines depict linear regression model fits between variables with 0.95 confidence intervals depicted by shaded areas. Spearman correlation coefficients and associated p-values are displayed in the upper left corners.

**Supplementary Figure 3:**
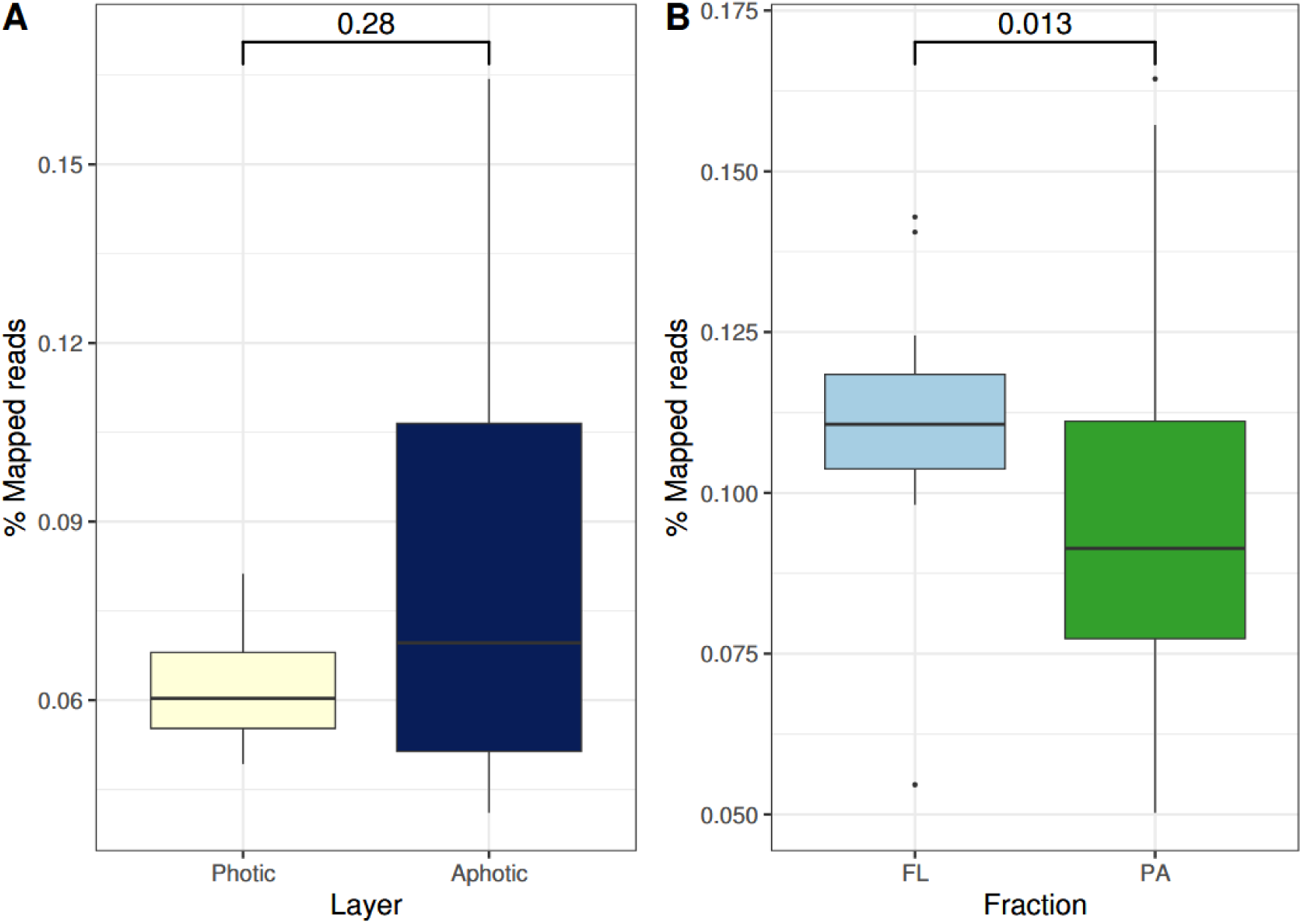
Boxplots depicting the percentage of total reads mapping to Cas proteins among metagenomes from samples grouped by photic zone (A) and grouped by size-fraction (B). Boxes depict the median, the first and third quartiles. Whiskers extend to 1.5 of the interquartile ranges. Outliers are represented as dots above or below whiskers. The *p-value* derived from Mann-Whitney tests comparing the significance of the difference between values of each group is shown above.

**Supplementary Figure 4:**
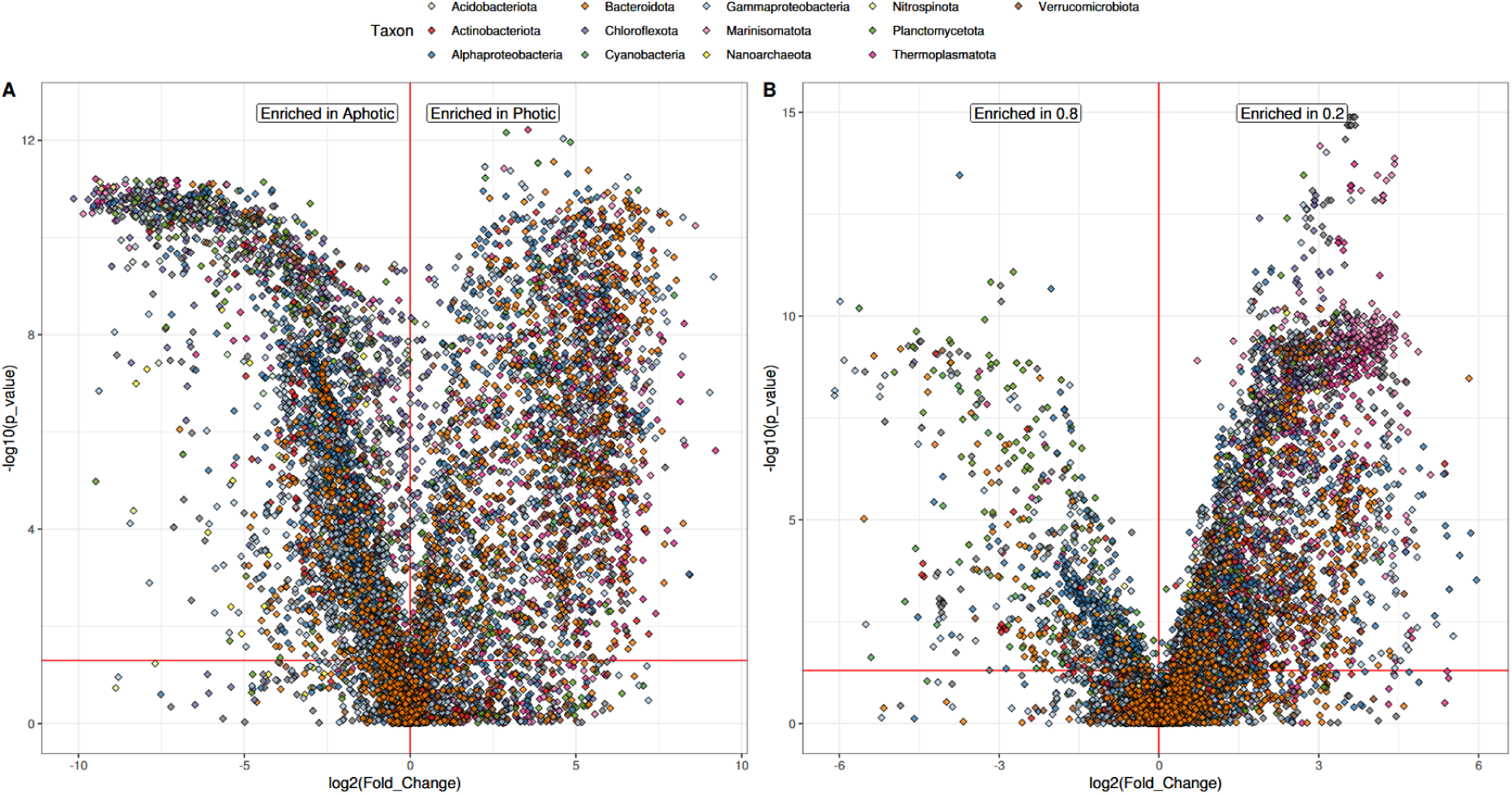
Volcano plots depicting the enrichment patterns of OceanDNA MAGs among Malaspina samples according to photic zone (A) and size-fraction (B). The *x*-axis depicts Log2(fold-change) of each MAG, where fold-change was calculated as mean abundance among samples from the right group divided by the mean abundance among samples in the left group. The *y*-axis depicts the -Log10(p-value) obtained from comparing the distribution of abundance values of the MAG among groups through the Mann-Whitney test. Each dot represents a different MAG and MAGs were coloured according to taxonomic affiliation. The horizontal red line depicts the 0.05 p-value threshold

**Supplementary Table 1:** Sample and Metagenome Metadata.

**Supplementary Table 2.** Annotation and classification of the inferred proteins belonging to Cas loci from the Malaspina Gene Catalog.

**Supplementary Table 3.** Number of predicted Class I and Class II Cas loci classified by types. The quality of the classification obtained through the majority rule method, and the presence of effector proteins as requirements for completeness.

**Supplementary Table 4:** Information on the 48 Cas9 proteins derived from the Malaspina Dataset.

**Supplementary Table 5:** Data availability. ENA accession numbers for all metagenomes in this study, including 43 newly released metagenomes from the nanoplankton (3-20 µm) size fraction.

**Supplementary File 1**: Fasta containing the sequences of 48 Cas9 proteins derived from the Malaspina Dataset.

## Notes

### Summary of Updates

New supplementary materials revision of the text

